# Lower cortical activation and altered functional connectivity characterize passive auditory spatial attention in ASD

**DOI:** 10.1101/2025.01.02.631088

**Authors:** Sergio Osorio, Jasmine Tan, Grace Levine, Seppo P Ahlfors, Steven Graham, Fahimeh Mamashli, Sheraz Khan, Robert M Joseph, Zein Nayal, Ainsley Losh, Stephanie Pawlyszyn, Nicole M McGuiggan, Matti S Hämäläinen, Hari Bharadwaj, Tal Kenet

## Abstract

Autism Spectrum Disorder (ASD) is a developmental condition characterized by difficulties in social interaction, communication and sensory processing. The ability to orient towards sounds is a key component of social interactions, yet auditory spatial attention remains relatively understudied in ASD, despite prior research indicating differences in this domain. Here, we investigate the neural signatures associated with passive auditory spatial attention in children with ASD (n = 21, ages 6-17) relative to age- and IQ-matched typically developing (TD) children (n = 31), using source-localized magnetoencephalography (MEG). Participants listened passively, while watching a silenced movie, to non-social auditory stimuli designed to either remain lateralized to one hemifield (*stay* trials), or to change in location from one side to the contralateral hemifield (*jump* trials). Linear mixed effects modeling showed lower cortical activation in the auditory cortex in the ASD group in response to *jump* trials, relative to the TD group. Additionally, functional connectivity analyses showed higher alpha-band functional connectivity in the ASD group between left auditory cortex seeds and right prefrontal and left parietal regions known to be recruited during auditory spatial attention. Right prefrontal alpha-band connectivity estimates were associated with behaviorally assessed auditory processing scores, whereas left parietal connectivity estimates were associated with ASD symptomatology. Our results align with the hypothesis that auditory spatial attention generally, and specifically orientation to sounds even when experienced passively, differs in ASD individuals.

## 1. Introduction

Autism Spectrum Disorder (ASD) is a developmental condition characterized by a complex set of behavioral and sensory symptoms, the core of which is differences in social communication (Ben-Sasson et al., 2019; Grzadzinski et al., 2013; Haesen et al., 2011; Hedger et al., 2020; O. Miguel et al., 2017). Among its sensory manifestations, differences in processing of auditory stimuli have been consistently documented among individuals with ASD (Alho et al., 2021; Alho, Khan, et al., 2023; Alho, Samuelsson, et al., 2023; Bharadwaj et al., 2022; DePape et al., 2012; Gonçalves & Monteiro, 2023; Haesen et al., 2011; Mamashli et al., 2017; Marco et al., 2011; O’Connor, 2012; Port et al., 2015, 2017; Schafer et al., 2020; Williams et al., 2021). Such differences observed in adolescence and adulthood might be the product of atypical neurodevelopmental trajectories, which have been previously observed when investigating auditory evoked potentials, such as the M50 and M100, in pediatric ASD populations (Edgar et al., 2020; Oram Cardy et al., 2004; Paetau et al., 1995; Ponton et al., 2002; Wunderlich et al., 2006).

Auditory spatial attention, i.e., our ability to perceive and track the location of sounds in space, is a complex phenomenon involving several stages of sensory and cognitive processing. For a sound to be successfully located in space, sensory encoding of acoustic features, specifically the relative timing and amplitude differences between ears, need to be integrated with efficient selective attention and object recognition processes (Fritz et al., 2007; Kayser et al., 2005; van der Heijden et al., 2019). This integration allows for environmentally adequate behavioral responses to be generated. In addition to primary and secondary auditory areas in the Superior Temporal Gyrus (STG), fMRI research has shown that locating a sound in space recruits a frontoparietal network of cortical regions (Degerman et al., 2006; Deouell et al., 2007; Haist et al., 2005; Kong et al., 2014; Smith et al., 2009; Zatorre et al., 1999). These regions include the ventrolateral Prefrontal Cortex (vlPFC), the dorsolateral Prefrontal Cortex (dlPFC), the precentral sulcus and gyrus, the central sulcus, the postcentral sulcus and gyrus, and the Inferior and Superior Temporal Lobes (IPL/SPL). Unsurprisingly, these areas overlap with well-defined functional networks observed in resting state fMRI data, including the ventral and dorsal attention networks, and the frontoparietal executive control network (Hames et al., 2016; Sadaghiani & Kleinschmidt, 2016; Supekar et al., 2013).

Research to date has shown that during auditory spatial attention tasks, individuals with ASD show poorer behavioral performance, measured as higher reaction times and reduced accuracy rates, when required to localize sounds in space (Soskey et al., 2017; Teder-Sälejärvi et al., 2005). While previous studies using electroencephalography (EEG) and magnetoencephalography (MEG) have established a range of differences in early sensory encoding of auditory stimuli in ASD (Bruneau et al., 1999; Jansson-Verkasalo et al., 2005; Orekhova et al., 2009; Roberts et al., 2010; Russo et al., 2009), these have been mostly documented in the context of general auditory processing. One study that directly investigated neural signatures in the context of auditory spatial attention in individuals with ASD showed that poorer behavioral performance in ASD individuals during auditory spatial attention tasks is accompanied by a lower amplitude of the N1 and the P3 components, two Event-Related Potentials (ERP) associated with early sensory encoding and attentional processing, correspondingly (Teder-Sälejärvi et al., 2005). Another study that also focused on auditory spatial attention in ASD found that individuals with an ASD diagnosis showed an absence of the Object-Related Negativity, an ERP component thought to reflect auditory segregation, as well as a reduced amplitude of the P400, an attention-dependent ERP response thought to reflect perceptual decision making (Lodhia et al., 2018). Notably, despite its importance for everyday life and communication, auditory spatial attention remains understudied in ASD.

To address this gap in the literature, we investigated cortical responses in ASD and Typically Developing (TD) individuals, ages 6 to 17, to direct manipulations of spatial lateralization cues in sound. Using source-localized MEG signals, we focused on evoked responses and functional connectivity from the auditory cortex to characterize the cortical signatures of differences in passive auditory spatial orienting in ASD. We also conducted correlation analyses to investigate possible associations between brain function and clinically relevant behavioral measures of ASD.

## 2. Methods

### 2.1. Participants

Thirty-one typically developing (TD, mean age = 13.1±3.4 years, range = 6-17) children, and twenty-one children diagnosed with autism spectrum disorder (ASD, mean age = 13.5±2.7 years, range = 7-17) participated in this study (Table 1). All participants met an IQ cutoff of full-scale IQ ≥ 70, as measured by the Kaufman Brief Intelligence Test II (KBIT, Kaufman & Kaufman, 2004). Participants were right-handed except for two ASD individuals, determined using the Dean Questionnaire (Dean, 1988). All participants had normal hearing, defined as pure tone thresholds better than or equal to 15 dB in both ears at octave frequencies between 250 Hz and 8 kHz, tested in a soundproof room using an audiometer. Informed consent was provided by parents or legal guardians, in addition to participants’ assent, according to protocols approved by the Institutional Review Board of the Massachusetts General Hospital.

**Table 1.** Characterization of participants. P-values are shown for independent samples t-tests for the difference of means across groups. NVIQ (non-verbal IQ), VIQ (Verbal IQ), ADOS (Autism Diagnostic Observation Schedule), SCQ (Social Communication Questionnaire), SRS (Social Responsiveness Scale), ASPS (Auditory Sensory Profile Score derived from the Sensory Profile Caregiver Questionnaire – see “Brain-Behavior Correlations below for description of the ASPS).

|  | ASD (N = 21, 4 females) |  |  | TD (N = 31, 4 females) |  |  | p-value |
| --- | --- | --- | --- | --- | --- | --- | --- |
|  | *N* | *Mean (SD)* | *Range* | *N* | *Mean (SD)* | *Range* |  |
| **Age** | 21 | 13.5 (2.7) | 7-17 | 31 | 13.1 (3.4) | 6-17 | 0.64 |
| **NVIQ** | 21 | 102.6 (16.9) | 59-136 | 20 | 113.7 (14.4) | 93-147 | 0.02 |
| **VIQ** | 21 | 101.2 (18.1) | 61-139 | 20 | 116.3 (12.6) | 96-144 | 0.001 |
| **ADOS** | 21 | 12.1 (3.7) | 7-18 | - | - | - | - |
| **SCQ** | 17 | 18.6 (7.3) | 8-31 | 22 | 4.4 (2.9) | 1-12 | < 0.001 |
| **SRS** | 21 | 78.6 (14.2) | 40-95 | 27 | 45.0 (6.3) | 34-61 | < 0.001 |
| **ASPS** | 20 | 3.1 (0.9) | 1.4-4.8 | 30 | 4.4 (0.5) | 3.4-5.0 | < 0.001 |

All ASD individuals had a formal clinical diagnosis of ASD and met criteria on the diagnostic Autism Diagnostic Observation Schedule, Version 2, (ADOS-2, Lord et al., 2000), administered by trained research personnel with demonstrated inter-rater reliability, the diagnostic Social Communication Questionnaire, Lifetime Version (SCQ Lifetime, Rutter et al., 2003), and quantitative Social Responsiveness Scale (SRS, (Constantino & Gruber, 2012)). All participants were also administered the Sensory Profile Caregiver Questionnaire (Dunn, 1999). ASD participants who met ADOS-2 criteria, but had a borderline ADOS-2 score, did not meet a cutoff of ≥ 15 on the SCQ, or a total t-score < 59 on the SRS, were further evaluated by an expert clinician (RMJ) to confirm the ASD diagnosis. A diagnosis of Fragile X syndrome, tuberous sclerosis, and other known risk factors, such as gestation for less than 36 weeks, were criteria for exclusion from participation in the study.

All TD participants were screened to have SCQ scores below the clinically relevant threshold. Parental report was used to rule out the possibility of any neurological or psychiatric condition (including psychotic episodes, seizures, clinical EEG abnormalities, Tourette syndrome, or any other tics), or substance abuse in the last six months. Participants with Tourette syndrome or tic disorders were excluded from both groups. Due to high comorbidity with ASD, participants with a diagnosis of ADHD, depression, anxiety or language delay were excluded from the TD group, but not from the ASD group.

### 2.2. Stimuli and procedures

The auditory stimuli consisted of 1200 ms broadband (0.5-2.5 kHz) noise bursts (Figure 1a). Stimuli were designed with a 400 microseconds Interaural Time Delay (ITD) between the stereo channels, thus generating a left- or right-lateralized percept depending on the direction of the ITD. In each trial, the broadband noise was amplitude modulated at either 25 Hz or 43 Hz (Ahlfors et al., 2024). To manipulate the ITD, discontinuities of ±400 µs were introduced in the noise carriers in each channel 550 ms after stimulus onset. Crucially, the discontinuities were designed to coincide with a trough in the amplitude modulation to render them monaurally imperceptible. In half the trials, the 400 µs discontinuities occurred in the same direction in both ears, thus holding the ITD constant for the full duration of the trial (Figure 1b). For the other half of the trials, carrier discontinuities were introduced in opposite directions, thus switching the ITD polarity and creating a percept of a spatial jump in the location of the sound from one hemifield to the contralateral side midway through the trial (Figure 1c).

**Figure 1.**
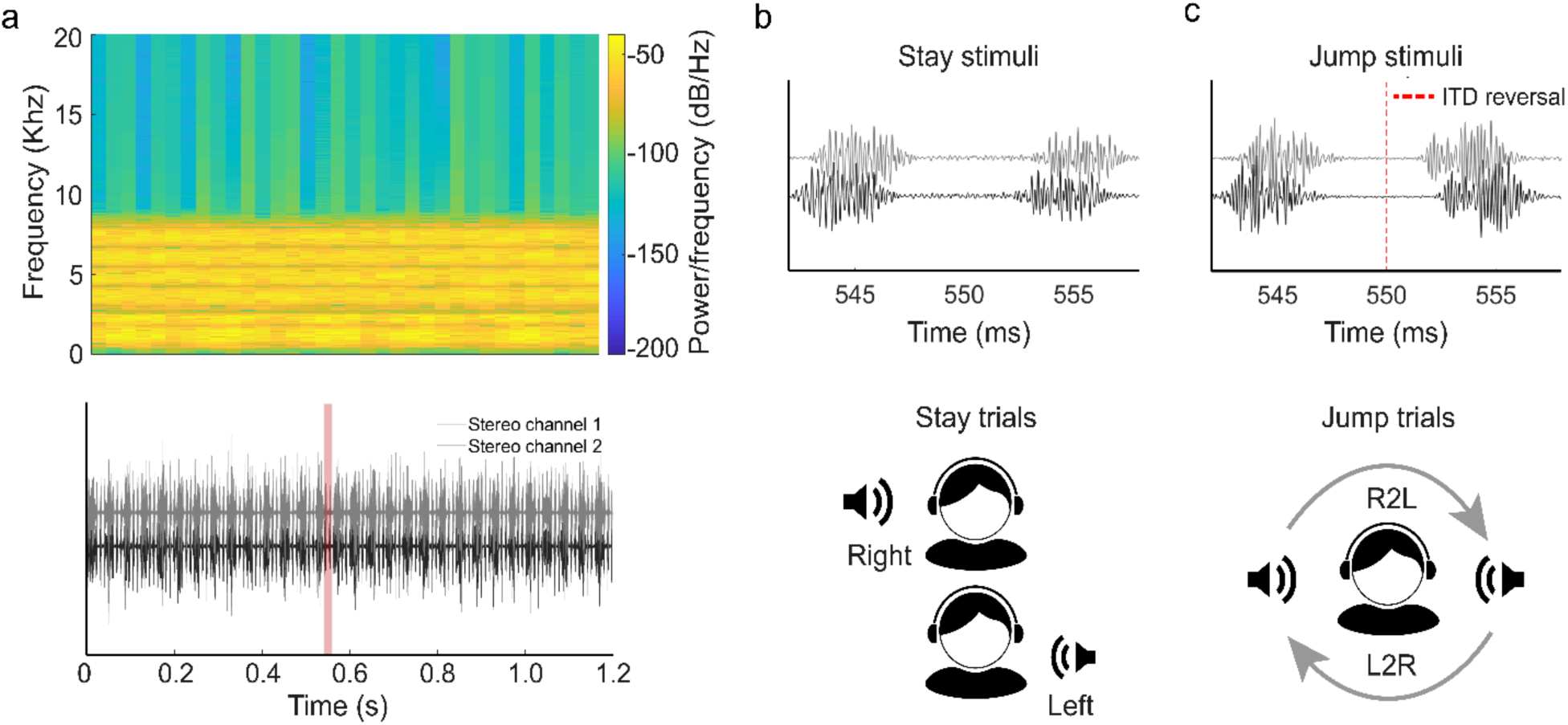
Stimuli and schematic representation of conditions. **a.** Spectrogram (top) and sound wave (bottom) of the auditory stimuli in the stay condition. The red shaded area in the bottom panel represents the time window from which panels in b and c (top) were extracted. **b.** For stay events, the directionality of the ITD was kept constant between the two stereo channels (top). Stimuli were presented with either left- or right-leading ITD (bottom), with equal probability. **c**. For jump events, the polarity of the ITD was reversed at 550 ms (dotted line, top). The sound was initially presented with either left- or right-leading ITD, with equal probability. The ITD reversal created a percept of a spatial jump in the lateralization of the sound, right-to-left (R2L) or left-to-right (L2R) (bottom).

The participants listened passively to the auditory stimuli while watching a silent movie with no subtitles. Participants were explicitly asked to ignore the auditory stimuli. In total, 200 trials were presented per condition. After rejection of bad epochs (see below for details), the mean total number of trials was 177 per condition, per participant. The mean number of trials across groups did not differ significantly (ASD jump = 182 ±20; ASD stay = 180 ±23; TD jump = 176 ±20; TD stay = 176 ±19). The interstimulus interval was uniformly distributed between 300 ms and 400 ms. Stimuli were presented at 65 dB-SPL via MEG-compatible earphones.

### 2.3. MEG Data acquisition

MEG data was acquired using a 306-channel Vector-View Magnetoencephalography system with 204 planar gradiometer and 102 magnetometer sensors (MEGIN Oy, Finland) inside a magnetically shielded room (IMEDCO, Switzerland). Signals were bandpass filtered between 0.1 and 330 Hz and sampled at the rate of 1000, or 3000 Hz and then downsampled to 1000 Hz. Head position was recorded during data acquisition using four head position indicator (HPI) coils attached to the scalp. The location of the HPI coils, the nasion, preauricular points, and additional points on the scalp surface were digitized using a Fastrack digitizer (Polhemus) system. The digitization information was used to co-register the MEG and MRI data for source modeling analyses. Electrocardiography (ECG) and electro-oculography (EOG) electrodes were used to detect heartbeat, saccades, and blinks. Five minutes of empty-room data were recorded at the end of the session to measure the instrumentation and environmental noise levels in the MEG sensors on the data acquisition date.

### 2.4. Structural MRI acquisition and processing

T1-weighted magnetization-prepared rapid gradient echo (MPRAGE) structural images were acquired using a 3T Siemens MRI scanner and a 32-channel head coil. Reconstruction of cortical surface, cortical volume and anatomical parcellation was conducted using FreeSurfer (Dale et al., 1999; Fischl, Sereno, & Dale, 1999). For group visualization of activation data, individual subjects’ cortical surfaces were co-registered using the spherical morphing method based on sulcal and gyral patterns (Fischl, Sereno, Tootell, et al., 1999).

### 2.5. MEG data preprocessing

MEG data was preprocessed and analyzed using MNE-Python (Gramfort et al., 2013). Noisy MEG channels were visually identified and excluded from further analyses. Temporal Signal Space Separation (tSSS, Taulu & Simola, 2006) was implemented to compensate for head movements and minimize noise from sources outside the virtual sphere circumscribing the MEG sensors. Next, data was bandpass filtered between 0.1 and 144 Hz, and notch filters were applied at 60 Hz and 120 Hz to remove line noise and its harmonics. Signal Space Projection (SSP, Uusitalo & Ilmoniemi, 1997) was used to remove cardiac and oculomotor artifacts by computing projections from the bandpass filtered (1-35 Hz) signal. The data were then epoched from −400 ms to 1400 ms with respect to sound onset and detrended to remove slow drifts. Epochs were excluded if the signal exceeded a peak-to-peak threshold of 4000 fT for magnetometer channels and 5000 fT/cm for planar gradiometer channels.

### 2.6. Source localization

Source localization was carried out by mapping the MEG data from the planar gradiometer sensors onto cortical space using MNE-Python (Gramfort et al., 2013). Source estimates were calculated on each participant’s individual anatomy, i.e., the FreeSurfer-reconstructed cortical surfaces from participant-specific T1 MRI scans. The forward solution was computed using a single-compartment boundary element model (BEM, (Hämäläinen & Sarvas, 1989)). Inner skull surface triangulations were generated from individual MRI structural data using the watershed algorithm. The cortical current distributions were obtained using the dynamic Statistical Parametric Mapping (dSPM, Dale et al., 2000) with a loose orientation constraint of 0.2 and depth weighting of 0.8 (Lin et al., 2006). To reduce the contribution of instrumentation and environmental artifacts to the MEG signals, a noise covariance matrix was obtained from empty room recordings. The noise covariance matrix was used to construct the inverse operator (Hämäläinen & Ilmoniemi, 1994) with a regularization parameter set to 0.1, and to noise-normalize the source estimates (Dale et al., 2000).

### 2.7. Auditory Regions of interest

For the source localization, it is important to use individual anatomy because MEG sensor space signals can show large interindividual variability due to the dependence on the location and orientation of the source currents within the convoluted cortical folding patterns (Hämäläinen et al., 1993). Thus, to reduce inter-subject variability in the group analysis, functional ROIs, rather than anatomically-based ROIs, were delineated for each participant when identifying early auditory ROIs. The auditory ROI for each participant was chosen to be the auditory area with greatest activation to the ITD reversal during *jump* trials, i.e. jump events. To determine the localization of this ROI in each participant, we first constrained the search by mapping onto the individual anatomy a search region (area outlined in black in Figure 2) that included portions of the following FreeSurfer labels: the Transverse Temporal Gyrus (TTG), the Transverse Temporal Sulcus (TTS), the Planum Temporale (PT), the Superior Temporal Gyrus (STG) and the Superior Temporal Sulcus (STS) (Destrieux et al., 2010). Note that since the STG label is very large, we restricted the search to the middle portion of the STG, corresponding to the section within the black contour in Figure 2. We then identified on each participant’s anatomical surface reconstruction the time-course of activations in response to jump events, using visual examination. The individual auditory ROIs were then chosen to correspond to the peak earliest activations following the jump event in the 575ms to 850ms time window from trial onset, i.e. 25ms to 300ms post jump event, and were manually delineated for each individual participant, in both the right and the left hemisphere.

**Figure 2.**
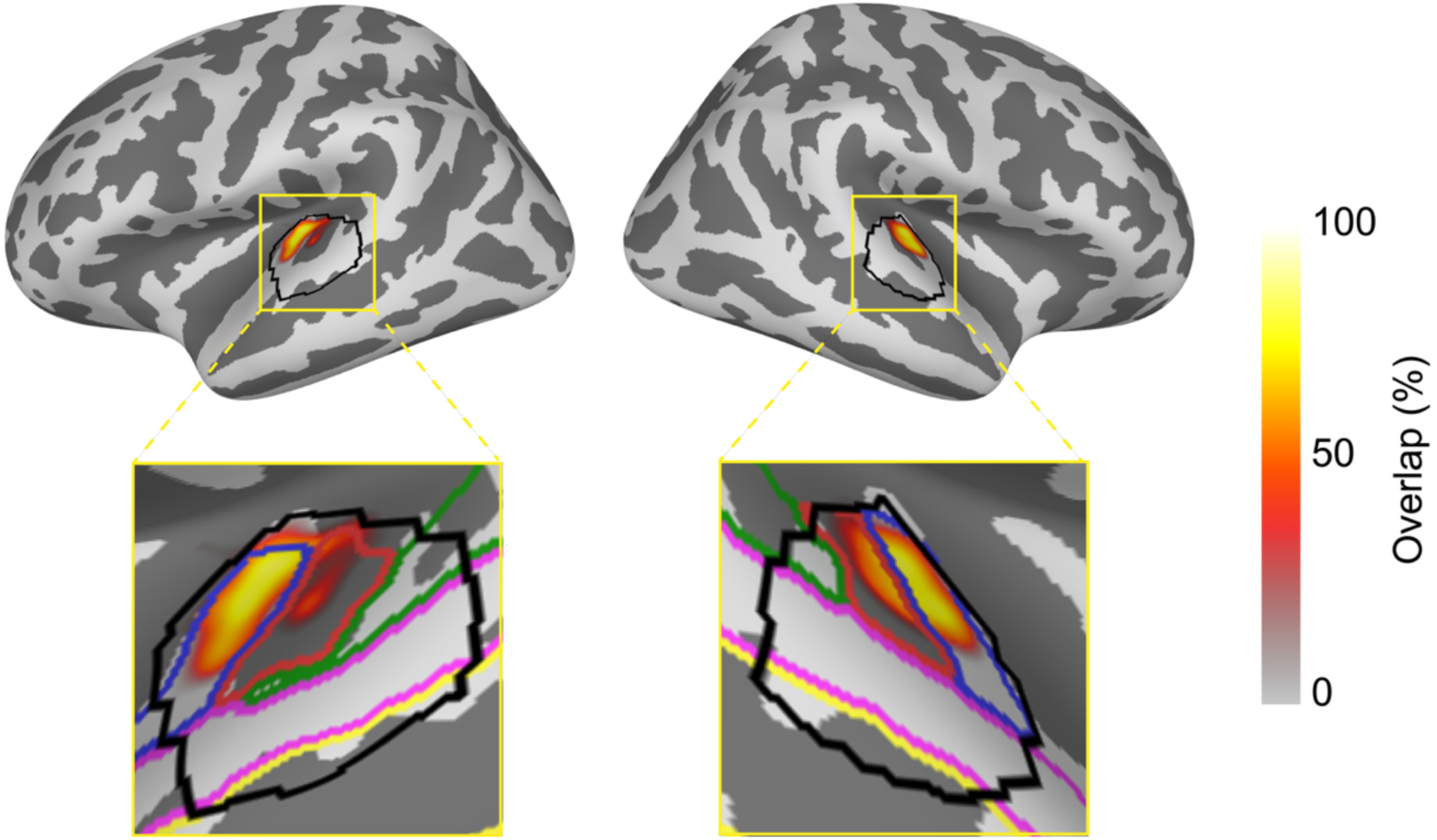
Extent of anatomical overlap across all participants for the manually drawn, left and right auditory ROIs. These ROIs were identified on the basis of cortical activations in response to ITD reversals in the Jumps condition, on the individual anatomy, and morphed onto common cortical space (Freesurfer’s fsaverage cortical surface) for visualization purposes. Zoomed-in windows show anatomical parcellations from the Destrieux atlas (blue: TTG, red: TTS, green: PT, magenta: STG, yellow: STS). The colorbar represents the extent of overlap in ROI across participants, i.e. the likelihood that a given spatial location was included in any participant’s ROI.

Figure 2 shows the overlap probability, meaning the spread of anatomical localization of the auditory ROI when mapped onto the fsaverage brain for visualization purposes, for left and right auditory ROIs, across all participants. ROI size was not significantly different across groups (ASD mean = 63 ±18 vertices, TD mean = 64 ±21, vertices; t = −0.21, p = 0.83).

### 2.8. Defining the response to jump events

The magnitude of the response to ITD reversal during *jump* trials, i.e. jump events, was defined as the area under the curve (AUC) between the onset of the response and the offset of the response to the jump event (AUCjump). The onset and offset latencies of the response were visually identified in each participant individually. To this end, we set the minimum onset time at 575 ms (i.e., 25 ms following the jump event), and the maximum offset time at 850 ms, and identified the onset, peak, and offset of the response within that time window individually for each participant. When responses were not easily identifiable, the full 575 to 850 ms time window was used to estimate the AUCjump for the response. There was no significant difference between the groups in either the onset latency (Left: TD mean = 671 ±46 ms, ASD mean = 670 ±35 ms, t(50) = 0.10, p = 0.92; Right: TD mean = 670 ±42 ms, ASD mean = 656 ±33 ms, t(50) = 1.26, p = 0.21) or in the width of the time window of response (Left: TD mean = 136 ±42 ms, ASD mean = 119 ±37 ms; t(50) = 1.51, p = 0.14; Right: TD mean = 147 ±47 ms, ASD mean = 135 ±40 ms, t(50) = 0.93, p = 0.36). The mean latencies of the onset and the width of the window in response to jump events for each participant are shown in Table S1 in the Supplementary materials.

### 2.9. Functional connectivity analyses

We computed seed-based functional connectivity for each condition (*stay* and *jump*) from the left and right auditory cortices to spatial auditory attention relevant regions of interest (ROIs) selected anatomically using FreeSurfer (Destrieux et al., 2010). The selected ROIs were the dorsolateral prefrontal cortex (dlPFC), the ventrolateral prefrontal cortex (vlPFC), central regions (precentral and central sulci, precentral and postcentral gyri), and parietal regions (postcentral and intraparietal sulci, superior and inferior parietal sulci), all bilaterally, using Freesurfer’s cortical parcellations (Destrieux et al., 2010), as shown in Table S2 and Figure S1 in the supplementary materials. Because the ROIs for functional connectivity were based on anatomical labels in fsaverage, we mapped the cortical space of each individual onto the fsaverage surface for these computations. To that end, we used the same co-registration of individual subjects’ cortical surfaces with the spherical morphing method based on sulcal and gyral patterns (Fischl, Sereno, Tootell, et al., 1999) as was used for visualization of activation at a group level. All functional connectivity computations were thus carried out in the fsaveraged brain. The seed time courses for functional connectivity analyses corresponded to the auditory cortical activations in response to *jump* events. Functional connectivity was quantified in each individual subject using the Phase Lag Index (PLI), a measure of phase synchronization that is insensitive to zero-lag estimates, thus effectively controlling for spurious connectivity due to field spread of MEG signals (Stam et al., 2007). To maximize power, we used the non-directional definition of PLI (with values ranging from 0 to 1), which combines leading and lagging phase differences.

Based on our prior studies where across-group differences were modulated by functional connectivity in the alpha (8-12 Hz) band (Alho, Khan, et al., 2023; Alho, Samuelsson, et al., 2023; Khan et al., 2013; Mamashli et al., 2021), here too we chose to focus on zPLI estimates specifically in the alpha frequency band. We focused on the time window between 550 and 850 ms, which corresponded to the onset-to-offset of the evoked response to the jump event. Data was resampled to 250 Hz, and the PLI was computed for each trial between the right and left seed ROI time course and the time course at each vertex, using the wavelet method (spanning four cycles) between 4 and 30 Hz, for a 2000-ms window between −400 and 1400 ms. PLI estimates in the jump condition were then normalized with respect to the *stay* condition to obtain a *z*-scored PLI value, hereafter referred to as zPLI (see supplementary materials). The zPLI estimate is a *z* value which reflects the extent to which connectivity during *jump* trials deviates with respect to the baseline condition (i.e. *stay*), while additionally controlling for the non-normal distribution of connectivity estimates and the difference in trial number across participants and conditions (Khan et al., 2013; Maris et al., 2007).

### 2.10. Cluster-based permutation statistics within anatomical ROIs

Clusters of vertices showing a significant difference in connectivity across groups within the target regions of interest were identified using cluster-based permutation statistics (Maris & Oostenveld, 2007). This test was conducted for the alpha band (8 – 12 Hz) by averaging functional connectivity estimates over the time dimension between 550 and 850 ms, a time window corresponding to the onset and offset of brain activation to *jump* trials. This statistical test was implemented for each seed (left and right) and target ROI (prefrontal, central and parietal) separately, using a one-way ANOVA with a cluster-forming threshold of F = 10.54 (corresponding to a p value = 1e-06) and 1000 permutations. Clusters were identified based on spatial adjacency. To achieve this, data from each participant was morphed to a FreeSurfer average cortical representation with 10,242 vertices per hemisphere (Fischl, Sereno, Tootell, et al., 1999). Cluster size was then estimated for each surviving cluster and results were filtered to keep only the top 5% biggest clusters. Finally, clusters were sorted by their statistical significance, which was obtained by summing F values at each vertex within the formed cluster, and only the most significant cluster per seed and anatomical ROI was considered for further analyses. As a post-hoc control analysis, we repeated the same functional connectivity analysis pipeline using Coherence instead of PLI, to ensure the robustness of the cluster identification approach. We found that the two ROIs that showed significant group difference with PLI also showed a significant group difference when Coherence was used as the connectivity metric, as shown in Figure S2 in Supplementary materials. Cluster-based permutation statistics were implemented using MNE-python (Gramfort et al., 2013).

### 2.11. Linear mixed effects modelling

Brain responses to *jump* trials were investigated using Linear mixed effects (LME) modeling with the LME4 package (Bates et al., 2015) in R version 2023.6.1. The predicted variable was the brain response to *jump* trials, quantified as the Area Under the Curve (AUC) at the activation post jump onset. The AUC for *stay* trials was obtained using the same participant-specific time window as for *jump* trials. Fixed effects included group (ASD, TD), condition (*stay*, *jump*), and hemisphere (left, right). To control for the within-subject nature of the task, a random effect for participant was included.

### 2.12. Brain-behavior correlations

Pearson’s correlation was estimated between normalized functional connectivity estimates extracted from the two surviving statistically significant ROIs that showed significant group differences (see Results) and the three quantitative behavioral measures: the ADOS score and the SRS score, both measures of Autism severity, and the Auditory Sensory Profile Score (ASPS), a behaviorally assessed measure of auditory sensory processing symptoms in ASD. The ASPS was quantified as the average of the first five questions of the auditory section of the Sensory Profile - Caregiver Questionnaire, which measures auditory processing atypicality in terms of sensitivity to and distraction by noise. These questions are categorically different from questions 6-8, which are related to social communication (e.g., ignoring what parents say or not responding to their name being called, question 6-7) and to enjoying or seeking to produce strange noises question 8), rather than to more basic auditory processing. One participant with ASD did not have an ASPS score and was therefore excluded from analyses involving this measure. Multiple comparisons were addressed by adjusting *p* values using the False Discovery Rate (FDR).

## 3. Results

### 3.1. Evoked responses in the auditory cortex

Both ASD and TD individuals showed stronger activations during *jump* trials compared to *stay* trials (i.e., a jump-evoked response). AUC values between onset and offset of peak activations during *jump* events for both conditions are presented in Table 2.

**Table 2.**
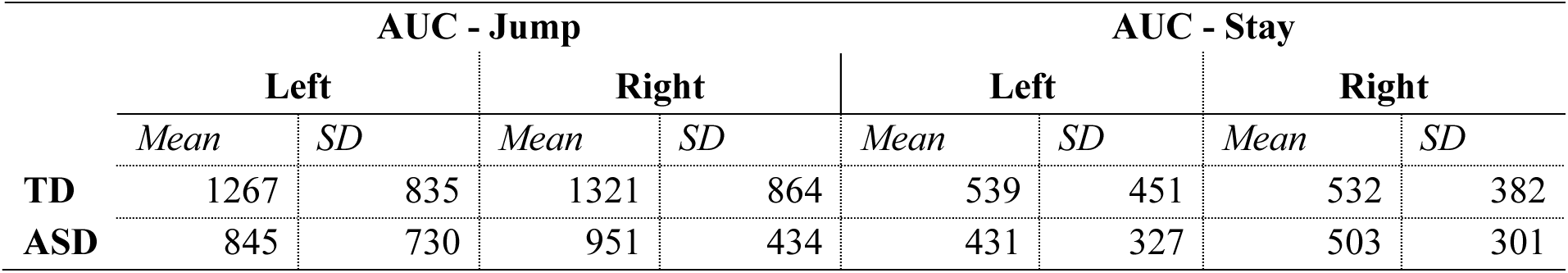
Area Under the Curve (AUC) for TD and ASD individuals at peak between onset and offset of brain activations to *jump* trials. The table shows the mean values by group and condition, as well as the standard deviation from the mean (SD).

|  | AUC - Jump |  |  |  | AUC - Stay |  |  |  |
| --- | --- | --- | --- | --- | --- | --- | --- | --- |
|  | Left |  | Right |  | Left |  | Right |  |
|  | *Mean* | *SD* | *Mean* | *SD* | *Mean* | *SD* | *Mean* | *SD* |
| **TD** | 1267 | 835 | 1321 | 864 | 539 | 451 | 532 | 382 |
| **ASD** | 845 | 730 | 951 | 434 | 431 | 327 | 503 | 301 |

Figure 3 shows the mean source time course for ASD and TD individuals in left and right auditory ROIs. Linear mixed effects modeling confirmed a significant effect of group (F (194.10) = 7.84, p = 0.006), condition (F (165.53) = 52.31, p = 1.67e-11) as well as a statistically significant interaction between group and condition (F (165.53) = 3.97, p = 0.048). There was no significant effect of hemisphere in predicting the AUC in brain activations (F (165.53) = 0.47, p = 0.49). Post-hoc pairwise comparisons showed that, within each group, mean AUC values for *jump* trials were significantly higher than *stay* trials (ASD: t (171) = 3.39, p = 9.0e-04, TD: t (171) = 7.26, p < 1.0e-4, insets in Figures 3a and 3b). Across groups, AUC values during *jump* trials were significantly higher for the TD group compared to the ASD group (t (185) = 3.39, p = 9.0e-05) in both left (Figure 3c) and right (Figure 3d) auditory ROIs. Maximum Likelihood Estimation analyses confirmed that the model outperformed a null (intercept-only) model in predicting evoked responses to jumps χ^2^(7) = 62.78, p = 4.19e-12, BIC = 3209.95, R^2^_conditional_ = 0.32, R^2^_marginal_ = 0.24).

**Figure 3.**
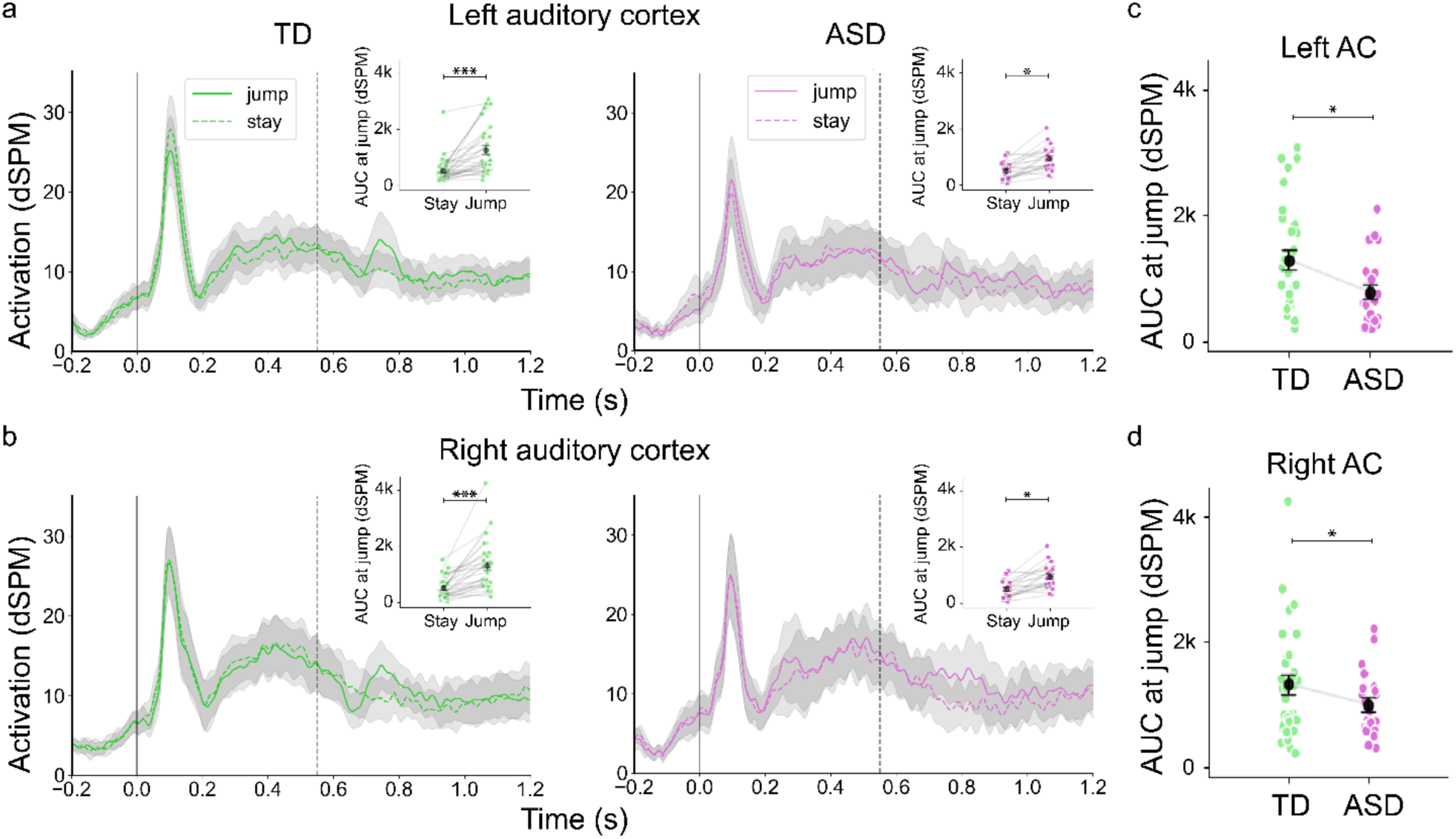
Evoked activations in auditory cortical ROIs. **a.** and **b.** Activations in the left (**a**) and right (**b**) auditory cortices for the TD (green) and ASD (purple) groups in response to jump (solid lines) and stay (dotted lines) trials. Solid vertical line at time zero represents sound onset. Dotted vertical line at 0.55 s represents the onset of and ITD reversal in the jump condition. Gray shades indicate the 95% confidence interval. Subpanels show the within group effect for stay and jump trials, by showing individual participant data changes across the two conditions. **c**. and **d**. Mean AUC values for *jump* trials for the TD (green) and ASD (purple) groups. Whiskers indicate the Standard Error of the Mean (SEM). *** p < 0.001, ** p < 0.01, * p < 0.05.

Activations in response to ITD reversals peaked on average at 734 ±52 ms in the left auditory cortex and at 724 ±48 ms in the right auditory cortex in the ASD group. In the TD group, the left hemisphere responses peaked at 742 ±50 ms, whereas the right hemisphere responses peaked at 739 ±51 ms. There was no statistically significant difference in peak latency across the two groups.

### 3.2. Functional connectivity between auditory areas and frontoparietal ROIs

Next, we investigated whether the spatial jump was also associated with different alpha-band functional connectivity dynamics, using zPLI, which corresponds to the PLI estimates for the *jump* condition normalized by PLI for the *stay* condition. Non-parametric cluster-based permutation tests showed two statistically significant clusters where alpha-band connectivity to the left auditory cortex differed between the two groups. The first cluster (n_vertices_ = 16, F = 44.29, p = 0.0010, Figure 4a, left panel) showed significant connectivity differences across groups between the left auditory seed and dorsolateral portions of the right prefrontal cortex (dlPFC). Within this cluster, alpha-band connectivity was significantly lower during *jump* trials compared to *stay* trials (median zPLI = −0.037 ±0.022) for TD individuals (t(30) = −10.28, p = 1.23e-10, Figure 4b, left), but higher during *jump* trials compared to *stay* trials (median zPLI = 0.021 ±0.027) for the ASD group (t (20) = −2.71, p = 0.015, Figure 4b, right). Additionally, prefrontal zPLI estimates significantly differed across groups (t(50) = −8.25, p = 1.23e-10, Figure 4b).

**Figure 4.**
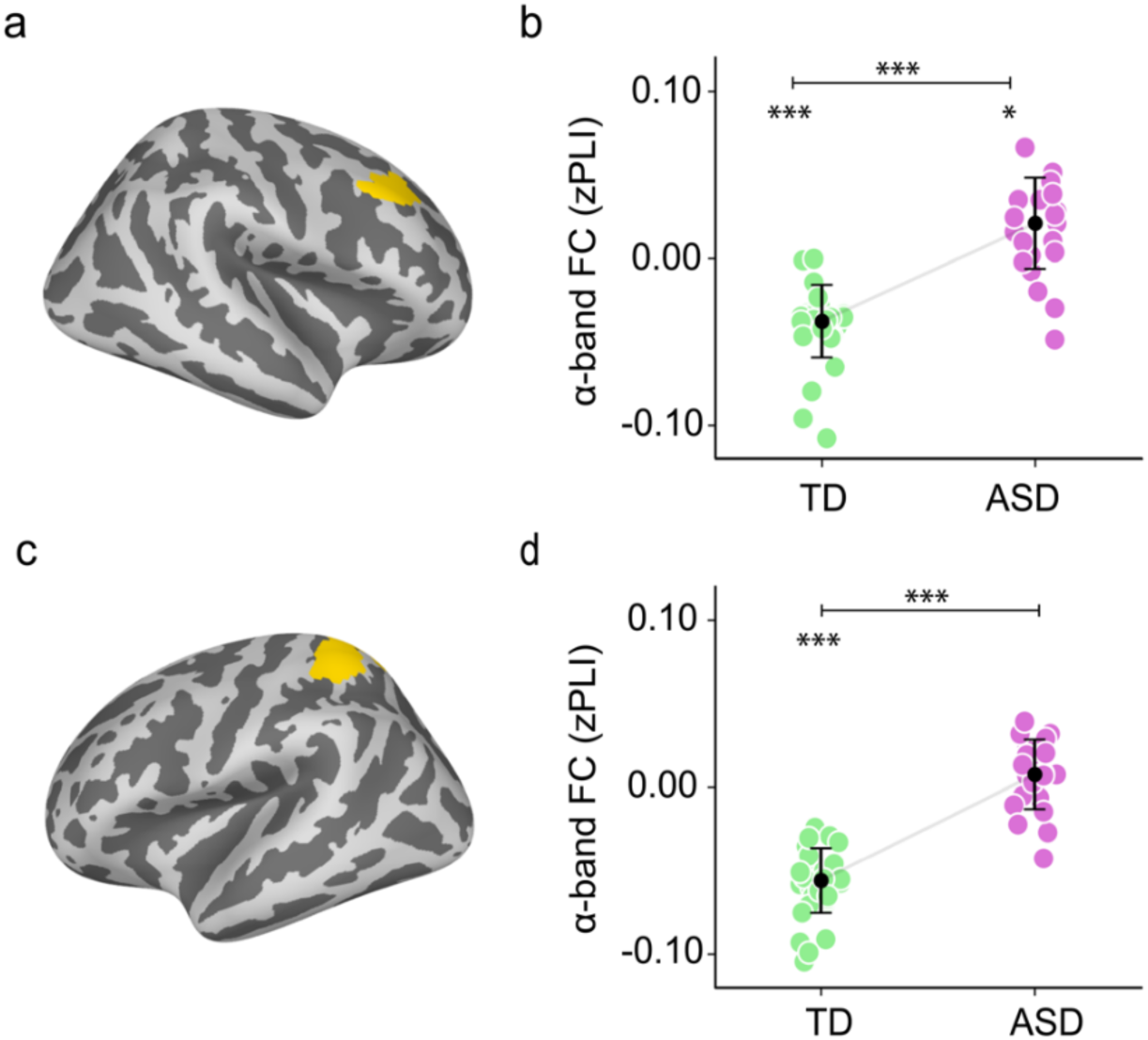
Functional connectivity with respect to the left auditory seed for the two statistically significant clusters during *jump* trials, normalized by *stay* trials. **a**. and **c**. Anatomical location of the statistically significant clusters in right prefrontal (**a**) and the left superior parietal regions (**c**). **b**. and **d**. Median alpha-band connectivity estimates obtained from the right dlPFC cluster (**b**), and left SPL cluster (**d**), for the TD (green) and ASD (purple) groups. Whiskers represent the Standard Error (SE). Asterisks on top of horizontal lines denote significant differences across groups. Asterisks below horizontal lines denote a significant difference against zero. *** p < 0.001, ** p < 0.01, * p < 0.05.

The second cluster (n_vertices_ = 17, F = 67.25, p = 0.0010, Figure 4c, left panel) showed significant connectivity differences across groups between the left auditory seed and portions of the left superior postcentral sulcus and superior parietal gyrus. Within this cluster, alpha-band connectivity was significantly lower during *jump* trials compared to *stay* trials (median zPLI = - 0.06 ±0.02) for TD individuals (t (30) = −16.38, p = 1.49e-15, Figure 4d, left), but zPLI estimates during *jump* trials did not significantly differ from the *stay* condition (t(20) = 1.34, p = 0.19) for the ASD group (median zPLI = 0.01 ±0.02, Figure 4d, right). Parietal functional connectivity within this parietal cluster was also significantly different across groups (t(50) = −11.19, p = 1.44e-14, Figure 4d). All reported *p* values from comparisons of MEG measures between the two groups are FDR-adjusted.

### 3.3. Correlations between brain measures and extent of ASD symptoms

To test whether the MEG measures were associated with relevant ASD traits, we correlated the magnitude of the evoked responses, as well as estimates of normalized functional connectivity (zPLI) with behaviorally assessed phenotypic measures of ASD severity (the ADOS and SRS t-scores) and auditory processing (the ASPS score). We found that higher alpha band functional connectivity in the left parietal cluster showed a positive correlation with autism severity, as measured by the ADOS composite score (r(19) = 0.51, p = 0.02, Figure S3 in Supplementary materials). Post-hoc analyses revealed that this effect was driven by the Social Communication and Interaction subcomponent of the ADOS (ADOS-sci: r(19) = 0.45, p = 0.04 uncorrected, q = 0.16), although this correlation did not survive multiple comparisons correction (Figure 5a). Prefrontal connectivity in the alpha band was also negatively correlated with the auditory processing component of the ASPS (r(18) = −0.63, p = 0.003, q = 0.023), as shown in Figure 5b. No significant correlation was found between functional connectivity and the SRS scores or any of the SRS subcomponent scores.

**Figure 5.**
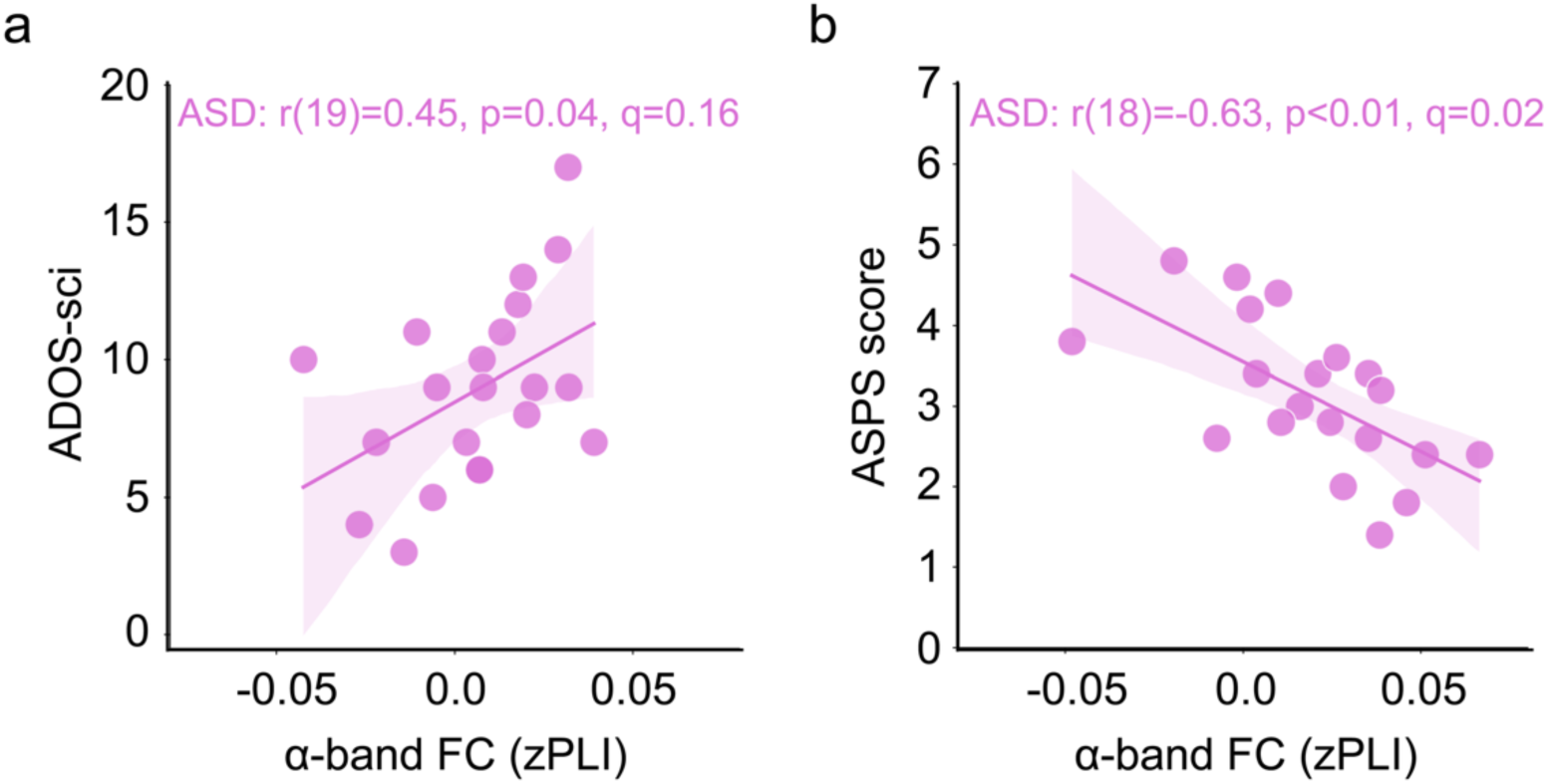
Brain-behavior correlations in the ASD group. **a**. Correlation between alpha-band connectivity estimates obtained from the parietal cluster and scores in the ADOS – Social Communication and Interaction subcomponent (ADOS-sci). (**b**) Correlation between alpha-band connectivity estimates obtained from the prefrontal cluster and the auditory processing section of the Sensory Profile – Caregiver questionnaire (ASPS score). Q values correspond to FDR corrected p values.

### 3.4. Correlations between brain measures and participants’ age

Given the evidence for different maturation trajectories in the M50 and M100 components of the auditory response in ASD, as discussed in Introduction, we also checked for possible effects of age. To that end, we computed post-hoc correlations between age and functional connectivity estimates, between age and the onset latency of the response to jump events, and between age and cortical activations. No significant correlations were found between age and any of the MEG measures of brain function.

## 4. Discussion

The aim of this study was to investigate passive auditory spatial attention in ASD. Specifically, we examined cortical activations and alpha-band functional connectivity in individuals with ASD and TD individuals in response to changes in sound location (i.e., *jump* events) under passive listening conditions. ASD individuals showed lower activation magnitudes in cortical auditory regions and altered alpha frequency band functional connectivity from left auditory areas to right prefrontal and left parietal areas. Additionally, functional connectivity estimates for the ASD group were significantly correlated with behaviorally assessed measures of ASD severity and auditory processing symptoms. Thus, our findings provide insight into the differential neural processing of spatial auditory information in ASD and TD groups.

### 4.1. ASD individuals show reduced auditory activation during jump trials

The fact that both groups showed cortical responses to *jump* trials confirms that both ASD and TD individuals were sensitive to acoustic cues indicative of a change in the location of a sound. This would be in line with findings which suggest that despite relative differences in performance, both groups can behaviorally locate sounds in space (Lodhia et al., 2018; Soskey et al., 2017; Teder-Sälejärvi et al., 2005). While both groups detected changes in localization, auditory cortical activations were significantly lower in the ASD group, which may be indicative of altered encoding of perceived acoustic features that are important for spatial mapping of auditory stimuli. These results are in line with previous electrophysiological research, which has shown a smaller amplitude of early and mid-latency auditory components during spatial (Lodhia et al., 2018; Teder-Sälejärvi et al., 2005) and non-spatial auditory processing tasks (Bruneau et al., 1999; Jansson-Verkasalo et al., 2005; Orekhova et al., 2009; Port et al., 2015; Russo et al., 2009; Soskey et al., 2017). They are also in line with postmortem human evidence in ASD of malformed brainstem circuits (i.e., superior olive) that are thought to be the first nuclei along the ascending auditory pathway to process binaural cues, such as interaural time delays (Kulesza et al., 2011). The significant group-by-condition interaction, where the ASD group exhibited a smaller magnitude of activation during *jump* trials, further supports the notion that individuals with ASD exhibit atypical auditory processing, potentially resulting from altered encoding or binding of relevant acoustic features (Bharadwaj et al., 2022; Gonçalves & Monteiro, 2023; Hames et al., 2016; Keehn et al., 2010, 2017; Lodhia et al., 2018; Marco et al., 2011).

### 4.2. ASD and TD groups show opposite connectivity patterns in response to jump trials

Our functional connectivity analyses revealed significant differences in temporo-frontal and temporo-parietal alpha-band connectivity across the ASD and TD groups, with the ASD group showing stronger connectivity during *jump* trials relative to the TD group. These prefrontal and parietal ROIs are both part of the ventral and dorsal attention networks and are functionally implicated in novelty processing, reorienting of attention, and more specifically, auditory spatial attention (Degerman et al., 2006; Deouell et al., 2007; Haist et al., 2005; Kong et al., 2014; Smith et al., 2009; Zatorre et al., 1999). These results suggest that the ASD group may lack the typical modulation of connectivity in response to sensory changes. This could reflect a disruption in the dynamic adjustment of attentional networks required for processing auditory spatial shifts, especially given that this is a passive paradigm, where attention was presumably directed away from the sounds. Atypical functional connectivity involving the auditory cortex, measured with EEG or MEG, has been documented in ASD by our group and others (Alho, Khan, et al., 2023; Alho, Samuelsson, et al., 2023; Demopoulos et al., 2024; Mamashli et al., 2017). Long-range alpha-band functional connectivity has been previously proposed as a neurophysiological marker of attentional modulation via coordination of temporal windows of neural excitability across cortical space (Klimesch et al., 2007; Palva & Palva, 2011; Sadaghiani & Kleinschmidt, 2016). Higher alpha-band connectivity in the ASD group could therefore be associated with atypical synchronization of attentional networks involved in sensory integration and cognitive control, which are often reported as dysregulated in ASD. The lower connectivity observed in the TD group could reflect a more typical response to the auditory shift, suggesting more efficient attentional allocation and neural processing. Although not related to functional connectivity, previous studies on auditory spatial attention in ASD have reported lower amplitude of mid-latency ERP components associated with attention and executive function, namely the P300 and P400 (Lodhia et al., 2018; Teder-Sälejärvi et al., 2005). This lends additional support to the idea that, on top of sensory encoding, attentional processes also differ in ASD during execution of auditory spatial attention tasks.

### 4.3. Alpha connectivity in ASD correlates with autism severity and behaviorally assessed auditory processing symptoms

We found that higher temporo-parietal alpha-band connectivity was positively correlated with higher manifestations of ASD symptoms as measured by the ADOS. Specifically, we found that this effect was driven by scores in the Social Communication and Interaction subcomponent of the ADOS questionnaire (ADOS-sci). Although this correlation did not survive correction for multiple comparisons, the trend is consistent with other studies showing a correlation between reduced functional connectivity of auditory cortex with both cerebral and cerebellar ROIs and increased ASD symptoms severity (Alho, Khan, et al., 2023; Alho, Samuelsson, et al., 2023; Mamashli et al., 2017; Sharda et al., 2018; Wilson et al., 2022).

Additionally, higher alpha-band connectivity between auditory and prefrontal regions was negatively correlated with behaviorally assessed auditory processing symptoms, as measured by the Auditory Sensory Profile Score (ASPS) in the ASD group. With higher scores on the ASPS indicating more pronounced auditory processing symptoms, these results are aligned with lower connectivity being linked with poorer integration of auditory spatial cues. This is also in line with previous work in our group, showing that increased connectivity between auditory and parietal regions is associated with poorer auditory processing skills, as measured by the ASPS (Alho, Khan, et al., 2023; Bharadwaj et al., 2022).

The functional relevance of both prefrontal and parietal regions in ASD has been previously established in the fMRI literature. Parietal regions have shown increased activation in ASD individuals compared to TD controls during social orienting (Greene et al., 2011) and visuospatial tasks (Damarla et al., 2010), whereas dorsolateral prefrontal regions are often reported to show either decreased activation (Luna et al., 2002) or increased inter-individual variability (Hawco et al., 2020) in ASD during spatial working memory tasks.

The finding of higher alpha-band functional connectivity in ASD during spatial attention tasks is in line with similar findings observed in resting state which propose alpha-band hyperconnectivity as a characteristic feature of ASD (Demopoulos et al., 2024; Orekhova et al., 2014; Wang et al., 2020). These findings highlight the potential impact of prefrontal and parietal regions on spatial auditory processing in ASD, while also underscoring the role of alpha in driving atypical functional connectivity in ASD. The underlying neural mechanisms that result in the alpha band being prominently featured in studies showing atypical functional connectivity in ASD are likely to be complex. While a full discussion of such putative mechanisms is beyond the scope of this study, alpha oscillations are likely to be the result of complex interactions across both inhibitory and excitatory processes (Bollimunta et al., 2008; Bruining et al., 2020; Lozano-Soldevilla, 2018; Lozano-Soldevilla et al., 2014; Nelson & Valakh, 2015), and thus atypicality in either or both of those domains could well underlie the observed group differences in alpha-band functional connectivity.

### 4.4. Limitations and future studies

The relatively small sample sizes of the ASD and TD groups limit the power of the findings, and while the findings align well with prior studies, future studies with larger cohorts will be important for further confirming the observed group differences in cortical activations and connectivity patterns, as well as to potentially uncover additional group differences. In terms of the age range of our participants, although our analyses revealed no association between age and either of our neurophysiological variables of interest, we cannot rule out the possibility that the study was not sufficiently powered to capture neurodevelopmental differences between the groups, especially given prior evidence of more general differences in maturation trajectories of auditory processing in ASD (Edgar et al., 2020; Oram Cardy et al., 2004; Paetau et al., 1995; Ponton et al., 2002; Wunderlich et al., 2006). Furthermore, because our results were obtained during passive listening conditions, we lack task-relevant behavioral correlates that allow us to rule out the possibility that participants with ASD actively engaged auditory attention differently from the TD participants. Future studies with an active spatial auditory attention task and behavioral assessments of spatial auditory attention will be needed to elucidate any behavioral differences between the groups, and links between these differences and cortical measures, and to explore in more detail the meaning of the correlations with social (ADOS-sci) and sensory (ASPS), and how these two domains of atypicalities may interact in ASD. Lastly, even though the auditory cortex ROIs were drawn in the individual cortical reconstructions, the spatial specificity of these MEG data is likely not sufficiently precise to draw conclusions about whether there were individual differences in functional localization or whether individual neuroanatomy may influence differences in activations. It is possible that even within the predefined FreeSurfer anatomical label, not the exact same functional region was selected as the auditory ROI for each participant. Open questions that remain therefore include the neuroanatomical specificity of the individual responses, which would be best addressed with high-resolution fMRI in combination with EEG or MEG.

## 5. Conclusion

The results presented here provide novel insights into the neural mechanisms underlying differences in auditory spatial attention in ASD. We found that ASD was associated with lower auditory cortical activations and altered alpha-band functional connectivity between auditory cortex and frontal and parietal regions, in response to shifts in the spatial location of sounds. The neurophysiological measures were correlated with measures of ASD traits and behaviorally assessed auditory processing symptoms in the ASD group, thus confirming their relevance to the ASD phenotype. Future directions include the evaluation of these metrics with an active, rather than passive, auditory spatial attention paradigm, and extending it to larger samples as well as to related conditions, so as to elucidate the specificity of the results to ASD.

