## Supplementary Material for "Lower cortical activation and altered functional connectivity characterize passive auditory spatial attention in ASD"

**This file contains:**

1. Description of zPLI, and of PLI variability.
2. Supplementary Tables S1, and S2
3. Supplementary Figures S1, S2 and S3

### 1. Z-normalization of functional connectivity estimates, and stability of the PLI measure across individuals and groups

Alpha-band functional connectivity estimates (PLI) were obtained between the mean activation at the left and right auditory labels and all vertices in the frontal, central, parietal and temporal lobes. Metrics of functional connectivity show a non-gaussian distribution and are highly sensitive to sample size differences, which can introduce bias in connectivity estimations (Maris & Fries, 2007; Sekihara et al., 2011). To address these issues, we estimated the zPLI, and adaptation of the z coherence estimation (Maris & Fries, 2007) previously used in work from our group (Khan et al. 2013). zPLI is defined as:

$$zPLI = \frac{\left(\tanh^{-1}\|PLI_1\,(f)\| - \left(1/(N_1-2)\right)\right) - \left(\tanh^{-1}\|PLI_2\,(f)\| - \left(1/(N_2-2)\right)\right)}{\sqrt{\left(1/(N_1-2)\right) + \left(1/(N_2-2)\right)}}$$

Where $N_1$ and $N_2$ represent the degrees of freedom in the jump and stay condition, respectively, $PLI_1$ are the connectivity estimates in the jump condition and $PLI_2$ the connectivity estimates in the baseline condition (stay). The sign of the *zPLI* denotes whether connectivity is higher (positive) or lower (negative) with respect to the baseline condition. PLI was estimated in each subject's individual anatomy, which was subsequently morphed to the FreeSurfer fsaverage cortical template, after which connectivity estimates were z normalized and group comparisons carried out.

To evaluate the stability of functional connectivity across individuals in the ASD group, we compared the inter-subject variability in PLI between the stay and jump conditions for each ROI pair. Levene's test was used due to its robustness to non-normal distributions. We did not find any significant difference in variance in the ASD group for the left auditory to right prefrontal cluster (SD stay =0.003, SD Jump = 0.002, $F(40) = 2.63$, $p = 0.11$), nor for the left auditory to left postcentral/parietal cluster (SD stay =0.002, SD Jump = 0.002, $F(40) = 0.07$, $p = 0.8$) in this post-hoc test. This suggests that, at least under our experimental design, PLI estimates are stable across both conditions and across ROIs.


1. Khan, S., Gramfort, A., Shetty, N. R., Kitzbichler, M. G., Ganesan, S., Moran, J. M., ... & Kenet, T. (2013). Local and long-range functional connectivity is reduced in concert in autism spectrum disorders. *Proceedings of the National Academy of Sciences*, *110*(8), 3107-3112.
2. Maris E, Schoffelen JM, Fries P (2007) Nonparametric statistical testing of coherence differences. J Neurosci Methods 163(1):161–175.
3. Sekihara K, Owen JP, Trisno S, Nagarajan SS (2011) Removal of spurious coherence in MEG source-space coherence analysis. IEEE Trans Biomed Eng 58(11):3121–3129.

**Table S1.** Individual onset latencies and width of response windows (in ms) for all participants, as described in section 2.8, sorted by age. As a reminder, the Jump events occur at 550ms post-trial onset. Note that age did not correlate with either the response onset latency (ASD: left $r(19)=0.12$, $p=0.60$; right $r(19)=0.29$, $p=0.20$; TD: left $r(31)=0.26$, $p=0.16$; right $r(31)=0.12$, $p=0.52$) or width of the response window (ASD: left $r(19)=-0.06$, $p = 0.80$; right $r(19)=-0.14$, $p=0.52$; TD: left $r(31)-0.20$, $p=0.28$, right $r(31)=-0.08$, $p=0.65$). There was also no difference between groups in either onset latency or width of the response window (Section 2.8). Note that onset latency could be driven by several factors, including individual arousal levels, attention, and stress due to the MEG environment, among other factors that we did not control for.

| **ASD** | | | | | **TD** | | | | |
|---|---|---|---|---|---|---|---|---|---|
| ***Age*** | ***Onset latency*** | | ***Width*** | | ***Age*** | ***Onset latency*** | | ***Width*** | |
| | Left | Right | Left | Right | | Left | Right | Left | Right |
| 7.8 | 729 | 600 | 153 | 112 | 6.7 | 578 | 572 | 99 | 98 |
| 9.3 | 727 | 712 | 83 | 181 | 8.6 | 674 | 734 | 117 | 143 |
| 9.6 | 636 | 663 | 89 | 114 | 8.7 | 648 | 603 | 103 | 119 |
| 10.3 | 607 | 610 | 85 | 102 | 8.9 | 712 | 754 | 209 | 138 |
| 11.6 | 695 | 700 | 164 | 158 | 9.2 | 698 | 682 | 144 | 79 |
| 11.8 | 712 | 669 | 124 | 203 | 9.2 | 621 | 616 | 110 | 186 |
| 12.2 | 692 | 675 | 113 | 96 | 9.3 | 745 | 732 | 101 | 80 |
| 12.4 | 667 | 645 | 158 | 86 | 9.7 | 692 | 588 | 155 | 200 |
| 12.4 | 630 | 657 | 61 | 198 | 10.5 | 583 | 700 | 111 | 148 |
| 13.1 | 652 | 681 | 102 | 124 | 10.8 | 700 | 669 | 178 | 254 |
| 13.6 | 675 | 664 | 116 | 226 | 10.8 | 700 | 683 | 133 | 226 |
| 14.0 | 709 | 695 | 147 | 96 | 10.8 | 689 | 703 | 209 | 113 |
| 14.6 | 678 | 672 | 85 | 147 | 11.3 | 583 | 629 | 117 | 95 |
| 15.2 | 669 | 658 | 73 | 130 | 11.9 | 624 | 647 | 268 | 229 |
| 15.3 | 616 | 593 | 68 | 127 | 12.0 | 678 | 689 | 215 | 198 |
| 16.2 | 672 | 686 | 104 | 87 | 12.2 | 659 | 642 | 108 | 86 |
| 16.3 | 661 | 650 | 133 | 107 | 14.1 | 641 | 678 | 128 | 181 |
| 16.6 | 724 | 598 | 152 | 78 | 14.5 | 648 | 667 | 114 | 157 |
| 16.9 | 652 | 672 | 155 | 164 | 15.0 | 633 | 644 | 152 | 195 |
| 17.0 | 639 | 636 | 102 | 103 | 15.2 | 692 | 700 | 141 | 192 |
| 17.4 | 633 | 644 | 124 | 212 | 15.7 | 687 | 691 | 114 | 144 |
| | | | | | 15.9 | 716 | 645 | 139 | 151 |
| | | | | | 15.9 | 765 | 658 | 79 | 119 |
| | | | | | 16.2 | 725 | 730 | 93 | 111 |
| | | | | | 16.9 | 675 | 712 | 150 | 102 |
| | | | | | 17.0 | 733 | 657 | 76 | 144 |
| | | | | | 17.4 | 640 | 671 | 113 | 87 |
| | | | | | 17.6 | 695 | 698 | 110 | 107 |
| | | | | | 17.7 | 692 | 695 | 124 | 130 |
| | | | | | 17.8 | 641 | 644 | 158 | 164 |
| | | | | | 18.0 | 647 | 650 | 161 | 169 |

**Table S2.** Specification of target ROIs for cluster-based permutation statistics. Labels correspond to anatomical parcellations in the Destrieux 2009 atlas. Number of vertices are presented for labels in the left and right hemisphere, for the downsampled fsaverage source space.

| | **Destrieux-2009 label** | **No. Vertices** | |
|---|---|---|---|
| | | **lh** | **rh** |
| **Prefrontal** | S_front_sup | 187 | 175 |
| | S_front_middle | 87 | 127 |
| | G_front_middle | 267 | 255 |
| | S_front_inf | 136 | 147 |
| | G_front_inf-Triangul | 61 | 67 |
| | G_front_inf-Opercular | 107 | 98 |
| | ***Total*** | ***845*** | ***869*** |
| **central** | S_precenrtal-inf-part | 121 | 117 |
| | S_precenrtal-sup-part | 114 | 122 |
| | G_precentral | 238 | 241 |
| | S_central | 121 | 117 |
| | G_postcentral | 311 | 301 |
| | ***Total*** | ***594*** | ***597*** |
| **Parietal** | S_postcentral | 277 | 261 |
| | G_parietal_sup | 263 | 209 |
| | S_intrapariet_and_P_trans | 277 | 269 |
| | G_pariet_inf-Angular | 190 | 205 |
| | G_pariet_inf-Supramarginal | 250 | 263 |
| | S_interim_prim-Jenssen | 29 | 49 |
| | ***Total*** | ***1286*** | ***1256*** |

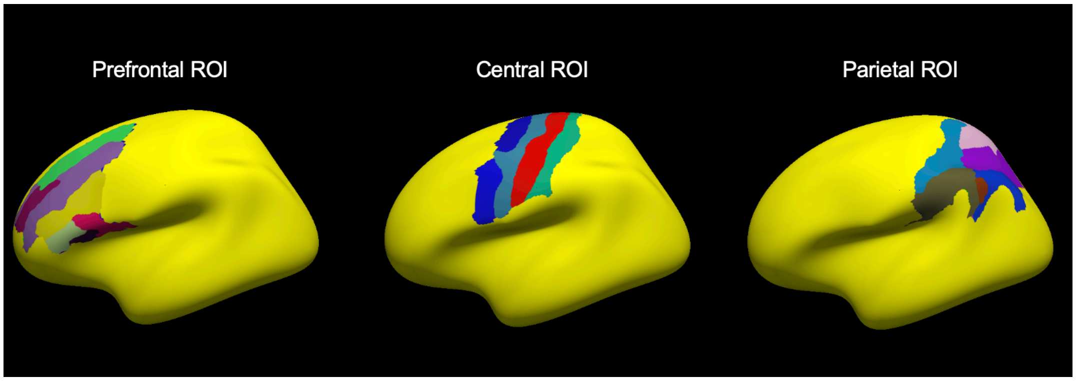


**Figure S1.** Regions of Interest (ROIs) relevant to spatial auditory attention, selected anatomically using FreeSurfer, here shown for the left hemisphere. Seed-based functional connectivity analyses were conducted from seed regions in the left and right auditory cortices to each one of these ROIs, bilaterally.

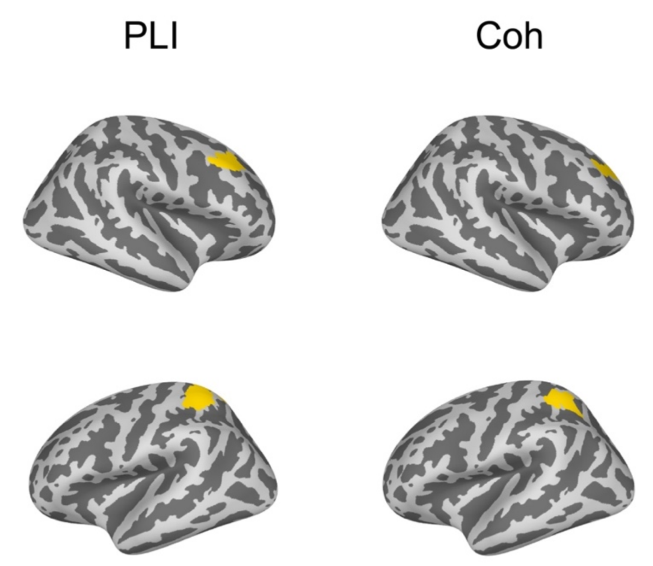


**Figure S2.** Clusters showing a statistically significant group difference across two functional connectivity methods: Phase Lag Index (PLI, left) and Coherence (right). We conducted post-hoc functional connectivity analyses using Coherence to assess the robustness of our results. Results show significant clusters of group differences using coherence that overlap with the two clusters reported here using the PLI.

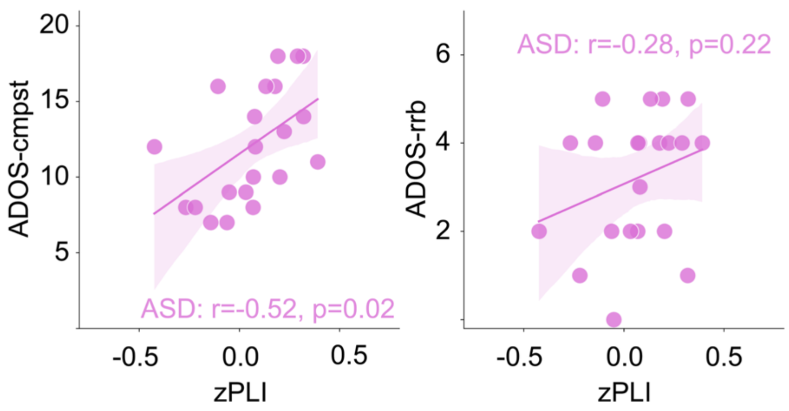


**Figure S3.** Correlations between functional connectivity estimates obtained from the superior parietal cluster and ADOS scores. The association was significant when functional connectivity data was correlated the ADOS composite score (ADOS-cmpst), shown in the left panel. Post-hoc analyses showed that this effect was driven by the Social Communication and Interaction subscore of the ADOS questionnaire (ADOS-sci, effect is reported in the main text), but the effect was not significant when correlated with the Restricted and Repetitive Behavior subcomponent (ADOS-rrb), shown in the right panel.
